# PP2A-B55α,δ phosphatase counteracts Ki67-dependent chromosome individualization during mitosis

**DOI:** 10.1101/2023.11.30.569375

**Authors:** María Sanz-Flores, Miguel Ruiz-Torres, Cristina Aguirre-Portolés, Aicha El Bakkali, Beatriz Salvador-Barberó, Carolina Villarroya-Beltri, Sagrario Ortega, Diego Megías, Daniel W. Gerlich, Mónica Álvarez-Fernández, Marcos Malumbres

**Author notes:** Correspondence to: M.A.F. and M.M. equal contribution (Miguel Ruiz-Torres, Cristina Aguirre-Portolés).

## Abstract

Cell cycle progression is regulated by the orderly balance between kinase and phosphatase activities. PP2A phosphatase holoenzymes containing the B55 family of regulatory B subunits (PP2A-B55) function as major CDK1-counteracting phosphatases during mitotic exit in mammals. However, the identification of the specific mitotic roles of these PP2A-B55 complexes has been hindered by the existence of multiple B55 isoforms. Here, through the generation of loss-of-function genetic mouse models for the two ubiquitous B55 isoforms (B55α and B55δ), we report that PP2A-B55 α/δ complexes display overlapping roles in controlling the dynamics of proper chromosome individualization and clustering during mitosis. In the absence of PP2A-B55α/δ activity, mitotic cells display increased chromosome individualization in the presence of enhanced phosphorylation and perichromosomal loading of Ki-67. These data provide experimental evidence for a new regulatory mechanism by which the balance between kinase and PP2A-B55 phosphatase activity controls the Ki-67-mediated spatial organization of the mass of chromosomes during mitosis.

## INTRODUCTION

Cell division requires an ordered series of phosphorylation/dephosphorylation events, which are regulated by both protein kinases and phosphatases. Entry into mitosis is triggered by the activation of Cyclin-dependent kinase 1 (CDK1), whose activity is controlled at multiple levels and regulated by several feedback loops ^1^. Once active, CDK1-cyclin complexes phosphorylate over 1000 proteins and cause the major rearrangements that mark mitotic entry. Additional mitotic kinases, such as Aurora A, Aurora B or Polo-like kinase 1 (PLK1) support mitotic entry and progression by phosphorylating additional substrates ^2^. Proteolytic degradation of the CDK1-activating subunit cyclin B1during the metaphase-to-anaphase transition triggers mitotic exit. This event is accompanied by the dephosphorylation of mitotic substrates through the ordered activation of specific phosphatase complexes ^3^. In budding yeast, the phosphatase Cdc14 plays a major role in counteracting Cdk-dependent phosphorylation and it is essential for mitotic exit. However, this function does not appear to be conserved in mammals ^4^, and several phosphatases, mostly PP2A and PP1, have been involved in mitotic exit in higher eukaryotes ^3,5^.

PP2A is the major serine and threonine phosphatase in eukaryotic cells. In cells PP2A can exist in two complex types: a dimeric core enzyme consisting of a catalytic C subunit and a scaffold A subunit; and a heterotrimeric holoenzyme composed by the core dimer (A subunit + C subunit) in complex with a regulatory B subunit, which provides substrate specificity and subcellular localization ^6^. In mammals, there are 4 different subfamilies of B regulatory subunits: B/B55, B’/ B56, B’’ and B’’’, each one with multiple isoforms and with some splice variants within isoforms, which allow the formation of about 80 different PP2A configurations ^6^. Among them, PP2A in complexes with B regulatory subfamily isoforms (PP2A-B55) appears to be the main CDK1 counteracting phosphatase during mitotic exit. Knockdown of PP2A-B55 complexes by RNA interference delays nuclear envelope reformation and post-mitotic reassembly of the Golgi apparatus in human cells ^7^, and impairs dephosphorylation of CDK phosphosites and mitotic exit in mouse cells ^8^. More recently, it has become evident that CDK1 antagonizing phosphatases are equally important for the dynamics of mitotic entry. The discovery of Greatwall or Microtubule-Associated Serine-Threonine Like kinase (MASTL) ^9^, and its function as a specific PP2A-B55 inhibitor ^10,11^, has further confirmed PP2A-B55 as a critical CDK-counteracting phosphatase, which must be downregulated to ensure the maintenance of a stable mitotic state ^12,13^. However, the specific roles of PP2A-B55 phosphatases during mitosis are still poorly understood. This is partially due to the existence of four different isoforms of the PP2A subfamily of B regulatory subunits in mammals (Β55α, B55β, B55γ and B558), encoded by 4 independent genes (*PPP2R2A-D*). Β55α and Β558 are expressed in most tissues, but expression of B55β and B55γ is almost restricted to brain ^14-17^. These four isoforms display high sequence similarity, with around 90% of identity at the aminoacidic level. Although some functions have been attributed to specific B55 isoforms, it is not currently clear to what extent they perform isoform-specific functions in the cell cycle.

By using loss-of-function genetic mouse models of the two ubiquitous B55 isoforms (Β55α and B558), we report here overlapping roles of these PP2A-B55 complexes for timely mitotic entry and exit. Importantly, our data point to a new mitotic role for PP2A-B55 in timely regulating chromosome individualization and clustering during mitosis by controlling the perichromosomal loading of the surfactant protein Ki-67.

## RESULTS

### Overlapping roles of PP2A-Β55α and PP2A-Β558 during mitotic entry and exit

To generate genetic loss-of-function mouse models of PP2A/B55 function, we focused on B55α and B55δ, the two ubiquitous isoforms of the mammalian B55 family (Figure 1A). We first generated a conditional knock-out (KO) mouse model for *Ppp2r2a*, the gene encoding B55α, in which exons 5-8 may be deleted upon expression of Cre recombinase (Figure. S1A, B). Crosses among heterozygous *Ppp2r2a*(+/−) mice resulted in no alive homozygous *Ppp2r2a*(−/−) animals (Figure 1B), indicating that *Ppp2r2a* is essential for mouse embryonic development, as recently reported ^18^. We also generated a constitutive KO mouse model for B55δ, by targeting exon 3 of the mouse *Ppp2r2d* gene (Figure S1C-E). Inter-crosses of *Ppp2r2d*(+/−) mice resulted in viable homozygous *Ppp2r2d*(−/−) animals at the expected mendelian ratios (Figure 1B). Depletion of B55 was confirmed at the protein level in both models by Western-blot analysis of mouse embryonic fibroblasts (MEFs) with isoform-specific antibodies (Figure 1C). Because the germline null mutation of *Ppp2r2a* resulted in embryonic lethality, we intercrossed *Ppp2r2a*(+/lox) mice to obtain *Ppp2r2a*(lox/lox) MEFs, which upon infection with adenoviruses expressing Cre recombinase generated *Ppp2r2a*(μ/M) cells (Figure 1D, E). In addition, we inter-crossed *Ppp2r2a*(lox/lox) mice with *Ppp2r2d*(−/−) to generate a double KO model. *Ppp2r2d*(−/−); *Ppp2r2a*(lox/lox) MEFs derived from these mice constitutively lack B55δ and, upon expression of Cre recombinase, become null for both genes [*Ppp2r2d*(−/−); *Ppp2r2a*(μ/M)] (Figure 1D, F).

**Figure 1.**
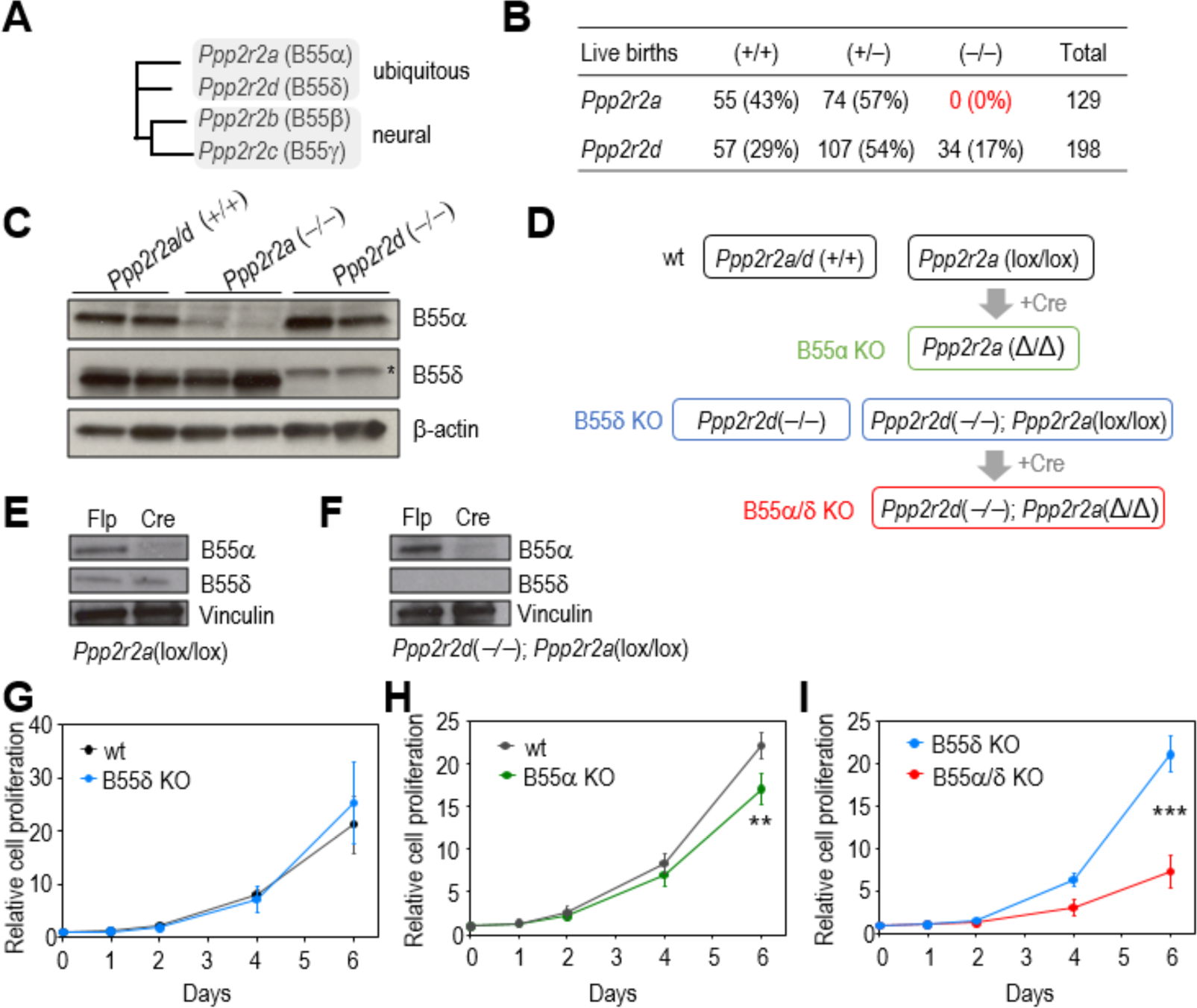
Effect of *Ppp2r2a* and *Ppp2r2d* deletion on mouse viability and cell proliferation. **A,** Phylogenetic tree of the four B55 mouse isoforms. **B,** Number (and percentage) of alive mice obtained with the indicated genotypes, from *Ppp2r2a*(+/−) and *Ppp2r2d*(+/−) heterozygous intercrosses, respectively. **C,** Western-blot analysis of MEFs with the indicated genotypes, using B55-isoform specific antibodies. Two clones of each genotype are shown. β-actin was used as a loading control. **D,** Schematic representation of the MEFs cell lines and genotypes used in this study. **E-F,** Transduction with Cre-expressing adenoviruses results in the depletion of *Ppp2r2a* compared to either control wt cells **(E)** or *Ppp2r2d*(−/−) cells **(F)** transduced with adenoviral vectors expressing Flp recombinases. Representative Western-blots for B55α and B558 in one clone of each genotype are shown. Vinculin was used as a loading control. **G** Relative cell proliferation of B558 KO [*Ppp2r2d*(−/−)] MEFs (n=3 clones) compared to wt cells (n=3 clones). **H,** Relative cell proliferation of B55α KO [*Ppp2r2a*(Δ/Δ)] cells, compared to control *Ppp2r2a*(lox/lox) cells (n=3 clones). **I,** Relative cell proliferation of B55α/δ KO [*Ppp2r2a*(Δ/Δ); *Ppp2r2d*(−/−)] compared to B558 KO [*Ppp2r2a*(lox/lox); *Ppp2r2d*(−/−)] cells (n=3 clones). In **G-I**, data are represented as mean ± SEM for each time point. \*\**p<*0.01; \*\*\**p*<0.001 (ANOVA).

We then analyzed the proliferation of these cells in which we eliminated either B55α, Β558 or both isoforms together. Whereas no significant differences in proliferation were observed between wild-type (wt) and Β558 KO cells (Figure 1H), a slight reduction in cell proliferation was detected in B55α KO cells (Figure 1H). Importantly, a severe impairment in proliferation was observed upon combined deletion of both isoforms, suggesting that both isoforms partially overlap in supporting cell proliferation (Figure 1I).

### PP2A-B55 is required for proper progression through mitosis

DNA content analysis of mutant MEFs did not show any major phenotype in single KO B55α or Β558 cells, but a small accumulation of cells with ≥4n DNA content was observed in Β55α/δ double KO model (Figure 2A and S2A). The balance between CDK activity and its counteracting phosphatase PP2A-B55 has been shown to be important both for mitotic entry and exit ^2,3^. To first explore the kinetics of mitotic entry upon PP2A-B55 depletion, cells stably expressing histone H2B fused to red fluorescent protein (RFP) were synchronized in G0 by serum starvation, and then stimulated with serum to re-enter the cell cycle. Entry into mitosis was monitored by time-lapse microscopy, and no differences were observed in B55α KO or Β558 KO compared with wt cells (Figure S2B, C). However, cells lacking both B55α and Β558, entered mitosis prematurely (Figure 2B and S2D). Despite this earlier entry into mitosis, the duration of mitosis (DOM) was prolonged in these Β55α/δ KO cells, but not in single KOs (80.97 ± 4.53 min in Β55α/δ KO vs 70.41 ± 4.25 min in Β558, 71.96 ± 13.92 min in B55α and 64.70 ± 12.41 min in wt cells) (Figure 2C). Interestingly, although no delay was observed from nuclear envelope breakdown (NEB) to metaphase, the timing from anaphase onset to nuclear envelope reformation (NER) was significantly increased in Β55α/δ KO cells compared to control cells (47.7 ± 3.14 min in Β55α/δ KO vs 34.73 ± 6.05 min in wt cells on average) (Figure 2C). In agreement with these observations, mitotic segregation defects, such as misaligned or lagging chromosomes, were observed in all mutant genotypes, being slightly more frequently in B55α compared to Β558KO, and much more frequently and severe in Β55α/δ KO cells (Figure 2D, E).

**Figure 2.**
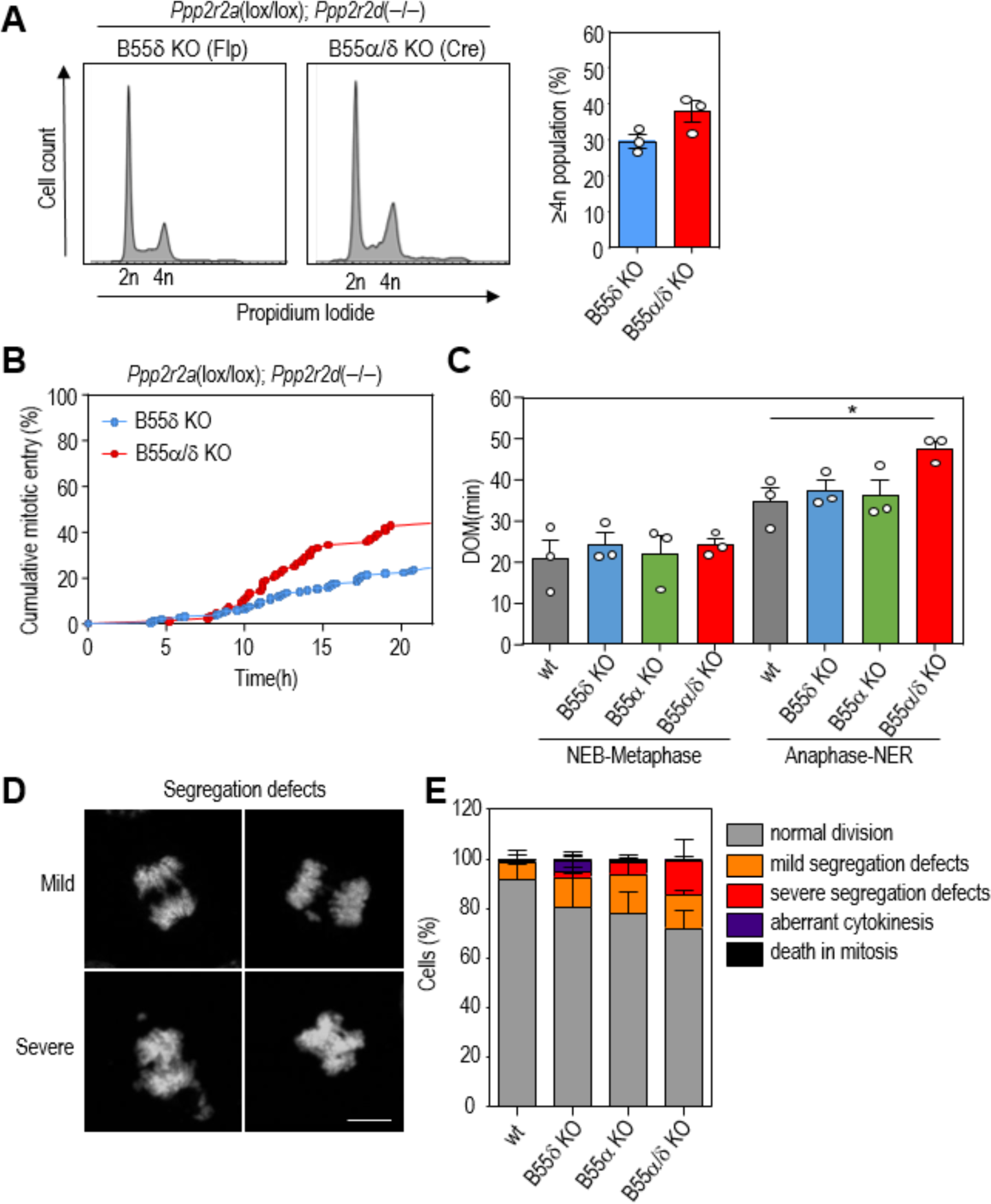
Mitotic defects in Β55α/δ deficient cells. **A,** Cell cycle profile (propidium iodide) of one representative *Ppp2r2a*(lox/lox); *Ppp2r2d*(−/−) clone 6 days after infection with AdenoFlp (B558 KO) or AdenoCre (B55δ KO) viruses. Bars on the right show the percentage of cells with ≥4n DNA content in 3 independent clones. **B,** Videomicroscopy analysis of mitotic entry (scored by cell rounding and chromosome condensation) in B55α/δ KO cells compared with single B55δ KO cells. **C,** Duration of mitosis (DOM) from NEB until metaphase; and, from anaphase onset until mitotic exit (flattening of daughter cells) in wt, B55α KO, B55δ KO and B55α/δ KO MEFs. **D**, Representative images (DAPI) of mitotic cells showing mild and severe chromosome segregation defects. Scale bar, 10µm. **E,** Quantification of mitotic defects detected in wt, B55α-deficient, B55δ-deficient and double KO B55α/δ- deficient MEFs. Data in C and E are means +SEM of n=3 different clones for each genotype, \**P*<0.05, 1-way ANOVA.

### Deletion of B55 induces chromosome scattering in the presence of nocodazole

To further investigate the defects observed during mitotic progression, we evaluated the robustness of the spindle assembly checkpoint (SAC) in B55-deficient cells, by performing time-lapse microscopy in the presence of the microtubule depolymerizing poison nocodazole. Both wt and B55-mutant cells displayed a significant arrest for about 9 ± 1h in prometaphase (Figure S3A). However, chromosomes were often scattered out of the main chromatin mass during the prometaphase arrest in B55-deficient cells (Figure 3A). The chromosome scattering phenotype was detected in about 27% of wt cells, more than 60% of Β55α and Β558 single KOs, and in almost all cells doubly deficient for both B55 isoforms (Figure 3B). Mitotic exit in the presence of nocodazole occurs through mitotic slippage, resulting in most cases in a single cell with one tetraploid nucleus or, in few cases, in one multinucleated cell in control cultures. (Figure 3C, D). Mononucleated cells after mitotic slippage were, however, observed only in about 25% of cells lacking either Β55α or Β558. In these single mutant cultures, about 50% of cells slipped out of mitosis as multinucleated cells, and the rest 25% died during mitosis, a cell fate that reached almost 80% in the combined depletion of Β55α and Β558 (Figure 3C and S3B). Given the high frequency of mitotic cell death in the Β55α/α double KO model, we focused on B55 single depletion models for further analysis of cell fate during mitotic slippage. In the presence of nocodazole, about 40% B55-deficient cells (41.43 ± 10.16 in B55α− and 35.75 ± 4.67 in Β558−deficient cells) exited mitosis as a single cell with a large number of small micronuclei (Figure 3E). Although micronuclei were also observed in about 20-25% wt cells, the number of micronuclei per cell was significantly higher in B55-mutant cells compared with wt cells (Figure 3F), in correlation with the presence of chromosome scattering in these mutant cells. Thus, B55 regulates the arrangement of mitotic chromosomes in a microtubule-independent manner.

**Figure 3.**
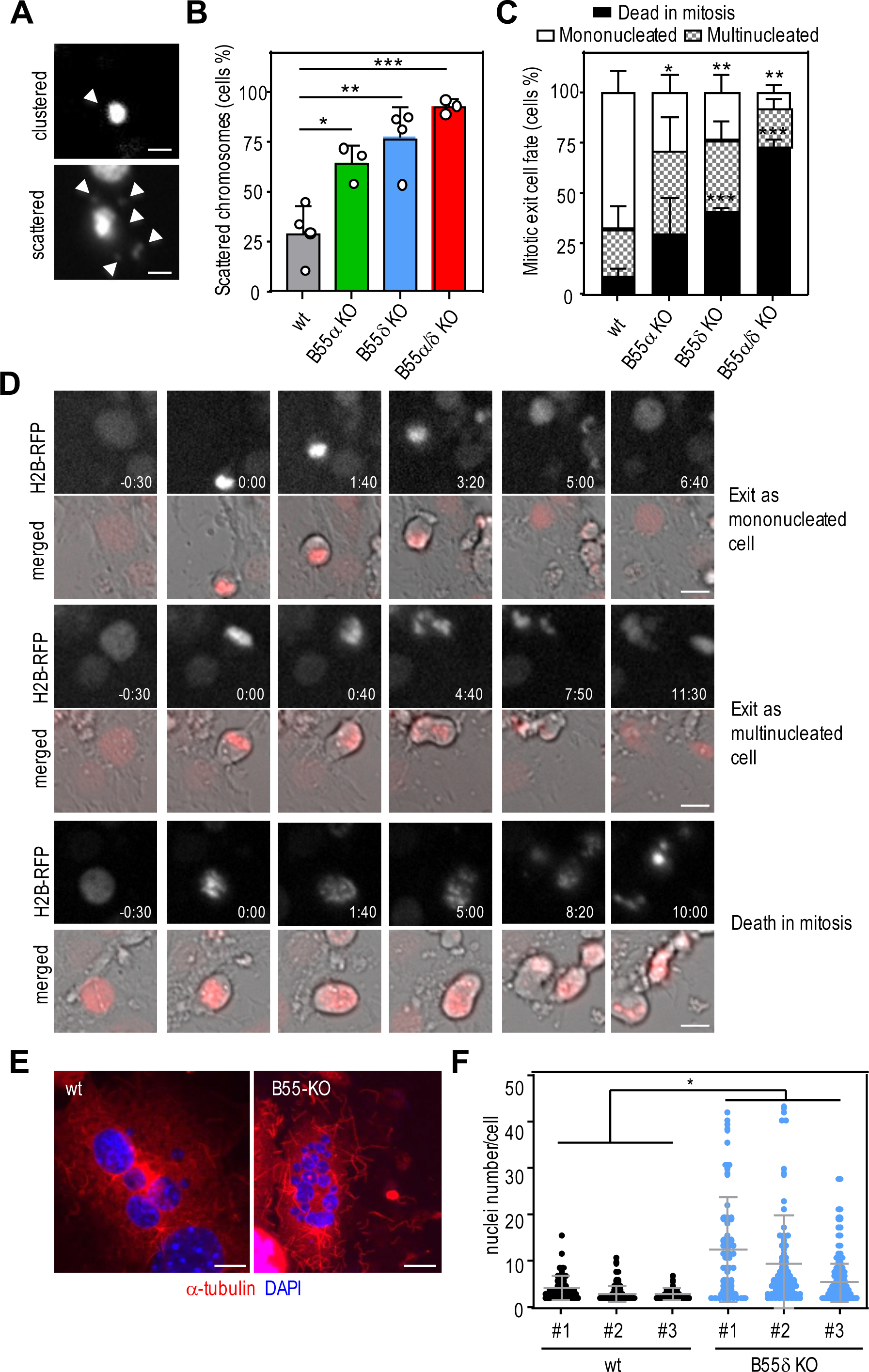
Chromosome scattering induced by deletion of B55 and nocodazole treatment. **A,** Representative images of H2B-RFP cells treated with nocodazole showing a clustered prometaphase (upper panel) and a prometaphase with scattered chromosomes indicated by white arrowheads. **B,** Percentage of prometaphase-arrested cells with scattered chromosomes in the indicated genotypes after nocodazole (0.8 µM) treatment. Data are means ± SEM of n=3/4 independent clones per genotype. \**P*<0.05, \*\**P*<0.01, 1-way ANOVA. **C,** Quantification of the mitotic exit cell fates (exit from mitosis as mononucleated cell, as multinucleated cell, or death during mitosis, see panel D for representative images) observed in the presence of nocodazole in each genotype; Data are means ± SEM of n=3/4 independent clones per genotype. \**P*<0.05, \*\**P*<0.01, \*\*\**P*<0.00, 2-way ANOVA **D,** Representative time-lapse images of H2B-RFP cells exiting from mitosis in the presence of nocodazole as a mononucleated cell (upper panel), as multinucleated cells (middle panel) or dying during the mitotic arrest (lower panel). **E,** Representative images of multinucleated cells stained with DAPI (blue) and α-tubulin (red) detected in wt and B55δ KO cells after nocodazole treatment. Scale bar: 10µm. **F**, Quantification of the number of nuclei per multinucleated cell in wt and B55δ KO cells after nocodazole treatment. Data are means ± SD of each clone (n=3/genotype). \**P*<0.05, Student’s t-test. Scale bars: 10 µm.

### PP2A-B55 regulates chromosome separation and Ki-67 accumulation at mitotic chromosomes

The chromosome scattering phenotype detected in B55-deficient cells might be due to repulsion between neighboring chromosomes during mitosis, which is regulated by the surfactant-like chromosome periphery protein Ki-67 ^19,20^. To quantify the degree of chromosome separation, we measured the area occupied by chromosomes in nocodazole-arrested prometaphase cells. Indeed, the area occupied by chromosomes was significantly higher in Β558 KO than in control cells, suggesting excessive separation of chromosome in the absence of B55 (Figure 4A). Prior work has shown that excessive separation between neighboring chromosomes can be induced by overexpressing Ki-67 ^20^ which led us to investigate whether the chromosome separation phenotype caused by B55 depletion might be due to misregulated Ki-67 levels on chromosomes. Indeed, Ki-67 accumulated to elevated levels in the perichromosomal layer in Β558 KO cells compared with wt cells (Figure 4B), suggesting that PP2A-B55 regulates Ki-67 levels on mitotic chromosomes.

**Figure 4.**
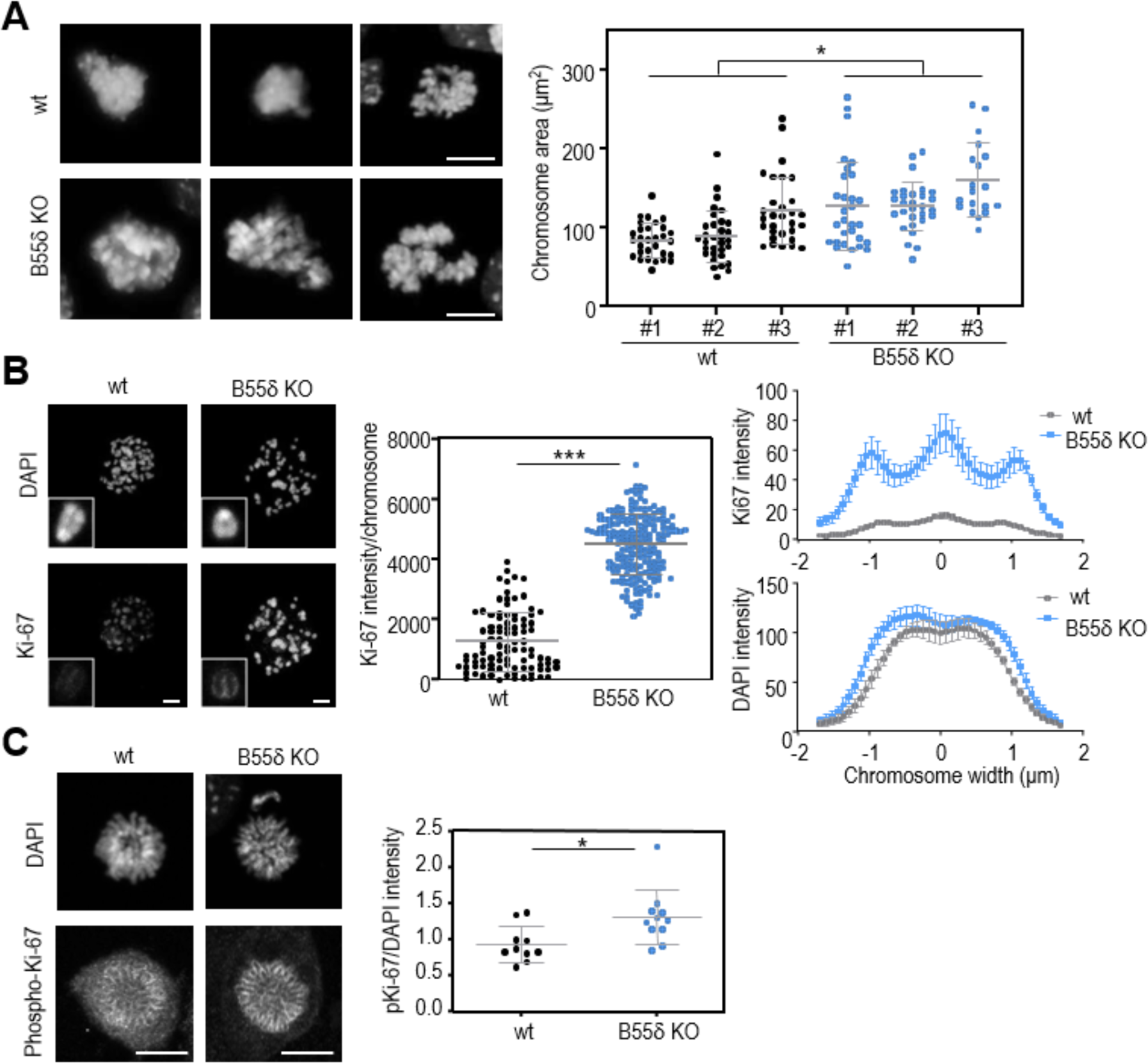
Effect of B55 depletion on chromosome separation and Ki-67 accumulation in prometaphase. **A,** Representative images (DAPI) of wt and B55δ KO nocodazole-arrested cells. The graph shows the quantification of the total chromosome area per cell. Data are means ± SD of each clone (n=3 per genotype). \**P*<0.05, Student’s t-test. **B,** Immunofluorescence images of metaphase spreads showing Ki-67 staining and DAPI in wt and B55δ KO cells. The graph shows the quantification of Ki-67 intensity per chromosome in at least 3 metaphase spreads/clone. \*\*\**P*<0.001, Student’s t-test. Line scan analysis of Ki-67 and DAPI intensity per chromosome width is also shown on the right. Data show the average intensity ± SEM of 3 metaphase spreads per clone (10 sections were analyzed in each prometaphase) **C,** Immunofluorescence images of prometaphases showing phospho-Ki-67 staining and DAPI in wt and B55δ KO cells. The graph shows the quantification of phospho-Ki-67 intensity normalized per DAPI in at least 10 cells per genotype. \**P*<0.001, Student’s t-test. Scale bars: 10 µm.

### Effects of B55 depletion on chromosome scattering are dependent on Ki-67

Ki-67 is hyperphosphorylated in mitosis at least partially by CDK1, and this phosphorylation is required for Ki-67 accumulation at the chromosome periphery ^21-24^. Ki-67 interacts and targets the phosphatase PP1 to anaphase chromosomes to revert mitotic histone phosphorylation during mitotic exit ^25^. However, PP1- binding is not required for its role in the assembly of the mitotic chromosome periphery, therefore, suggesting the involvement of other phosphatases. Interestingly, Ki-67 has also been identified as a putative substrate of PP2A-B55 complexes ^26,27^. In agreement with these observations, we detected increased levels of phosphorylated Ki-67 on condensed chromosomes in the absence of Β558 (Figure 4C), suggesting that the chromosome scattering phenotype of PP2A-B55 depletion might be caused through Ki-67 as a potential PP2A substrate.

We assessed whether Ki-67 is involved in the chromosome scattering phenotype induced by B55 depletion by transfecting Β558 KO MEFs with a siRNA against Ki-67 (Figure 5A,B). Importantly, depletion of Ki-67 suppressed the chromosome scattering phenotype in Β558KO cells to almost control levels (Figure 5C), and significantly reduced the fraction of cells exiting mitosis in the presence of nocodazole as multinucleated cells (Figure 5D). Thus, B55 regulates chromosome individualization during mitosis through the modulation of Ki-67 levels at the chromosome periphery.

**Figure 5.**
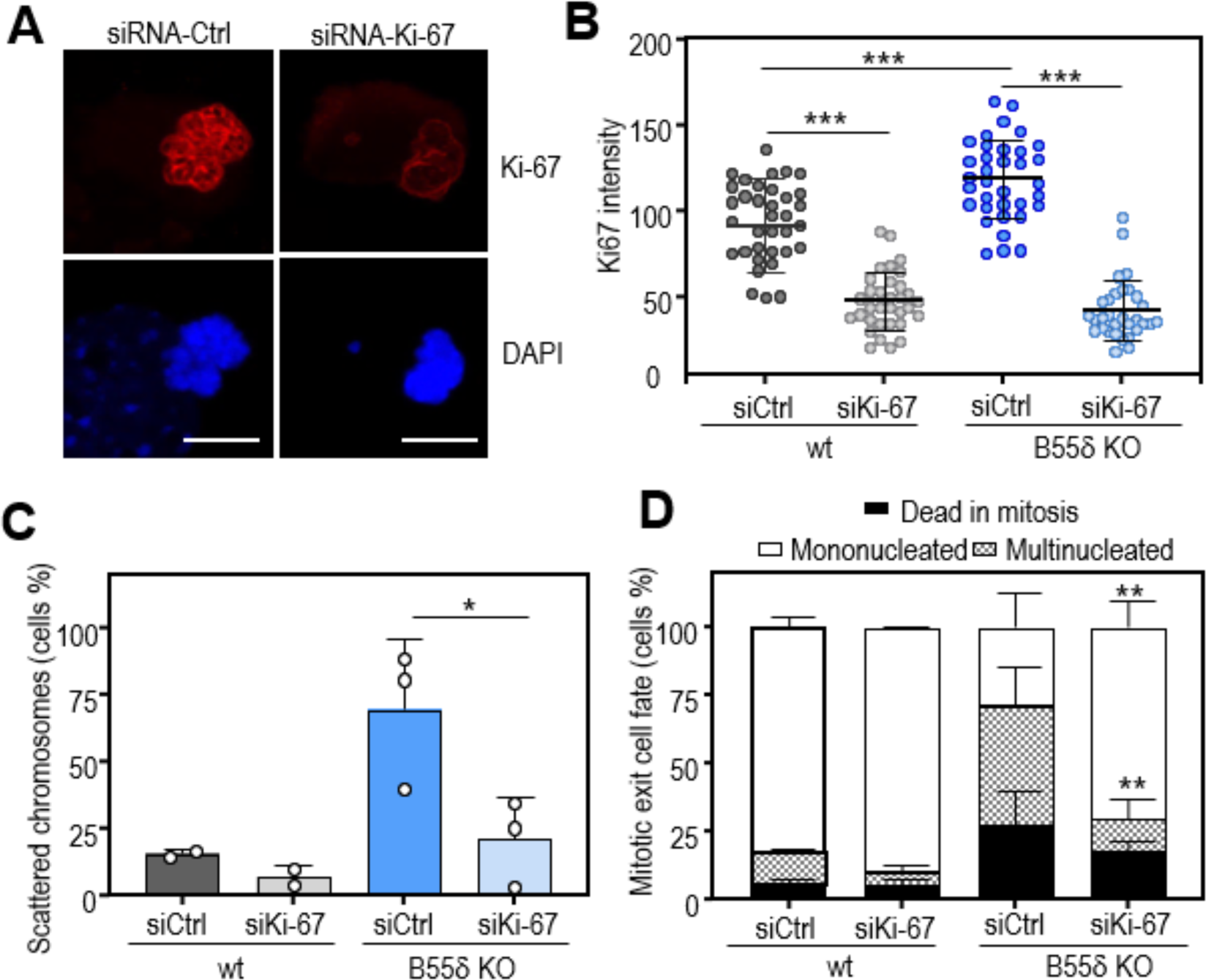
The chromosome scattering phenotype of B55 KO cells is Ki-67 dependent. **A,** Representative immunofluorescence images of wt MEFs transfected with a siRNA ctrl or a siRNA against Ki-67. Ki-67 (red) and DAPI (blues) stainings are shown. Scale bar, 10 µm**. B,** Quantification of the total Ki-67 intensity per cell in nocodazole-arrested prometaphases in wt and B55δ KO cells transfected either with siRNA control or siRNA against Ki-67. Data are means ± SD of one representative clone per genotype. \*\*\**P*<0.001, 1-way ANOVA **C,** Percentage of prometaphase-arrested cells with scattered chromosomes in the indicated genotypes after siRNA transfection and nocodazole (0.8 µM) treatment. Data are mean + SEM of n=2/3 independent clones per genotype. \**P*<0.05, Student’s t-test. **D,** Quantification of the mitotic exit cell fates (exit from mitosis as mononucleated cell, as multinucleated cell, or death during mitosis) observed in the presence of nocodazole in the indicated genotypes after siRNA transfection. Data are means +SEM of n=2/3 independent clones/ genotype. \*\**P*<0.01, 2-way ANOVA.

### PP2A-B55 regulates chromosome clustering in mitosis in human cells

To explore whether B55 also regulates the arrangement of mitotic chromosomes in human cells we next transfected Hela cells with siRNAs against human *PPP2R2A* (B55α) and *PPP2R2D* (B558) subunits (Figure 6A). Upon nocodazole treatment we also detected chromosome scattering, although to a lesser extent than in mouse fibroblasts. Importantly, both B55α and Β558 single-depleted cells, as well as Β55α/δ double-deficient ones, showed an increased frequency of this phenotype compared to scramble-transfected control cells, which was more significant in Β558 single knock-down (Figure 6B). In contrast to MEFs, mitotic slippage did not result in mitotic exit as multinucleated cells, and most nocodazole-arrested Hela cells either died in mitosis or exited as tetraploid mononucleated cells to die in the next interphase (Figure S4). To quantify the degree of chromosome clustering, we also measured the area occupied by chromosomes in nocodazole-arrested prometaphase Hela cells upon B55 depletion. In agreement with our data in MEFs, chromosome area was significantly higher in Β558 deficient cells compared to control cells (Figure 6C). Moreover, Ki-67 accumulation was also increased in Β558 depleted cells compared to control ones, suggesting a similar phenotype to the above described in MEFs (Figure 6D). Altogether, these data suggest that PP2A-B55 phosphatases contribute to proper chromosome distance during mitosis by regulating Ki-67 levels at the chromosome surface.

**Figure 6.**
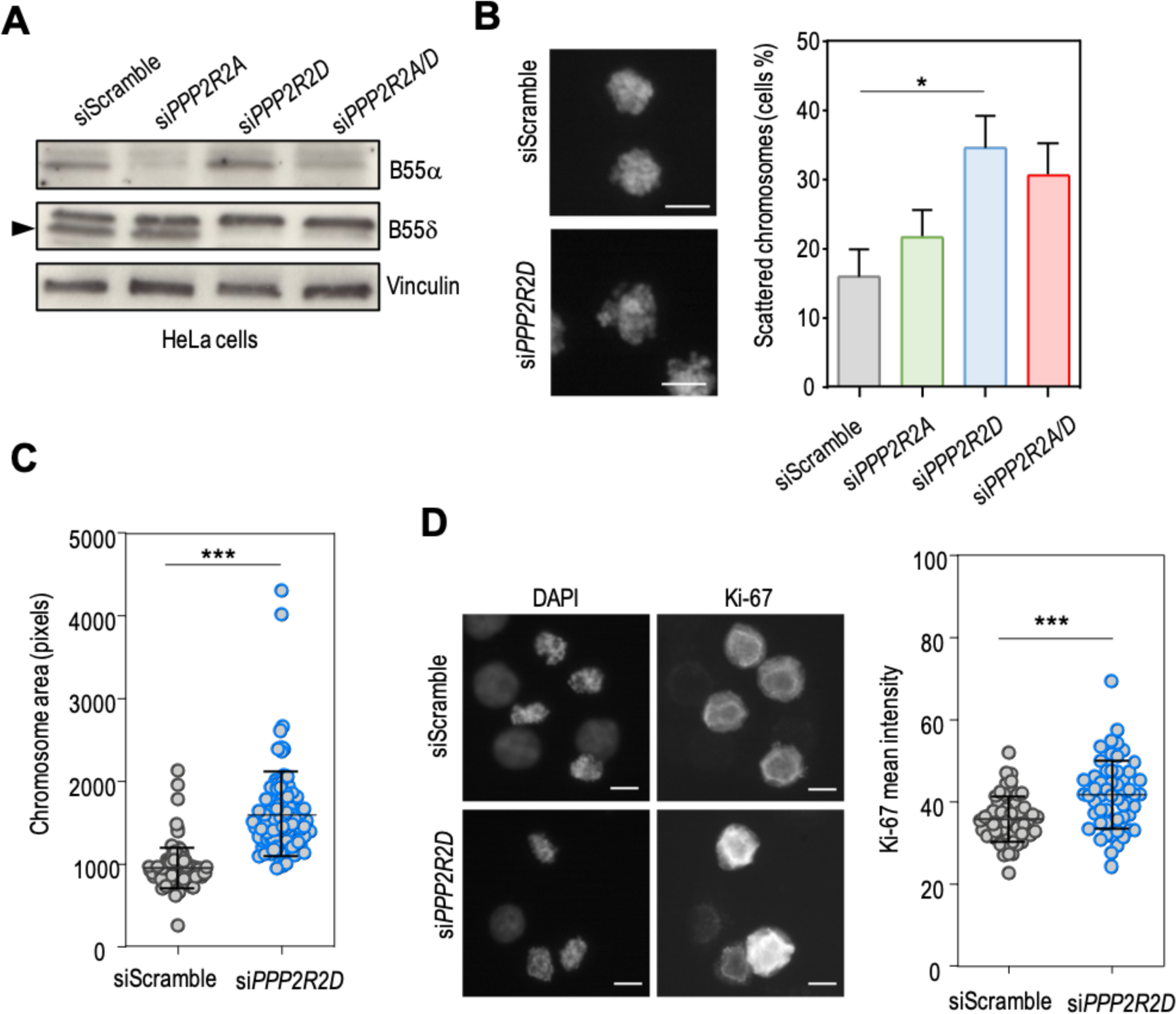
Effect of B55 depletion on chromosome individualization in nocodazole-treated Hela cells. **A**, Western-blot analysis of Hela cells transfected with the indicated siRNAs, using B55-isoform specific antibodies. Vinculin was used as a loading control. **B,** Representative images (DAPI) of control (siScramble) and B558-depleted (si*PPP2R2D*) nocodazole-arrested cells. The graph shows the percentage of prometaphase-arrested cells with scattered chromosomes upon transfection with the indicated siRNAs and nocodazole (0.8 µM) treatment for 14h. Data are means ± SEM. \**P*<0.05, 1-way ANOVA. Scale bar, 10 µm. **C,** Quantification of the total chromosome area per cell. Data are means ± SD of one representative experiment. \*\*\**P*<0.001, Student’s t-test. **D,** Representative immunofluorescence images of Hela cells transfected with a siRNA ctrl or a siRNA against *PPP2R2D*. Ki-67 and DAPI stainings are shown. The graph shows the quantification of the mean Ki-67 intensity per cell in nocodazole-arrested prometaphases. Data are means ± SD of one representative experiment. \*\*\**P*<0.001, Student’s t-test. Scale bar, 10 µm.

## DISCUSSION

The role of PP2A-B55 phosphatases in mammalian mitosis has mostly been studied in the context of mitotic entry or exit. Although single depletion of Β55α or Β558 does not affect the kinetics of entry into mitosis, combined depletion of both isoforms results in premature mitotic entry, indicating redundant roles for these two subunits in this mitotic function. This could explain why this phenotype has not been detected in previous genetic studies, in which single depletions were evaluated ^7,28^. In contrast, depletion of Β558 is sufficient to induce premature mitotic entry in *Xenopus* extracts, probably due to the low expression of other isoforms in this cellular system ^29,30^. The requirement of PP2A for mitotic entry seems to be contradictory with our previous data on MASTL (Greatwall) being dispensable for mitotic entry in mouse cells ^12^. However, it has recently been reported that PP2A-B55 is also directly inhibited by CDK1-mediated phosphorylation of its catalytic subunit to promote mitotic entry in a MASTL independent manner ^31^, which could explain the requirement of PP2A but not of MASTL for this function. A similar redundancy between B55 isoforms is also detected in the timing of mitotic exit, since only combined deletion of both subunits results in a significant delay from anaphase onset until nuclear envelope reformation, in agreement with our previous data showing the requirement of both isoforms for dephosphorylation of CDK mitotic substrates ^8^. In contrast, single depletion of Β55α was sufficient to delay mitotic exit in Hela cells ^7^, with no effect of Β558. These differences could be due to isoform-specific functions or to the differential expression levels of those isoforms between cellular systems. Indeed, Β55α appears to be the predominant isoform in many frequently-used human cell lines, whereas both Β55α and B55δ are both similarly expressed in mouse cell lines, at least at the mRNA level (unpublished data). Thus, the different degree of redundancy detected between isoforms is thought to reflect variations in co-expression levels across different organisms and cellular systems.

Our data also show for the first time the requirement of PP2A-B55 phosphatases for proper chromosome segregation during mitosis. While deletion of Β55α or B55δ only induce relatively mild missegregation, combined depletion of both isoforms causes prominent missegregation. This mitotic function also seems to be conserved since anaphase defects in chromosome segregation have already been described in two mutants of Twins, the only B55 isoform present in *Drosophila* ^32,33^. The severe segregation defects that we observed in the double B55α/δ KO cells might be the consequence of a premature mitotic entry in these cells, possibly before completion of DNA replication thereby preventing complete separation of sister chromatids during anaphase. Segregation defects might also be the result of a direct role of PP2A-B55 complexes in chromatid segregation. Interestingly, PP2A-B55 phosphatase have been related with centriole assembly and duplication in other organisms, such as *C. elegans* ^34^ and *Drosophila* ^35^, and, more recently, with centrosome maintenance in prostate cancer cells ^36^; and it is well known that centrosome dynamics alterations during mitosis may also lead to chromosome mis-segregation ^37^. Considering the wide spectrum of identified CDK mitotic substrates ^23,38^, a significant number of putative substrates of PP2A-B55 could participate in the regulation of all these processes, and further studies will be required to identify the most relevant ones in each mitotic phase.

Our data also reveal a novel function of PP2A-B55 phosphatase complexes in the regulation of chromosome separation during mitosis. In contrast to the previously discussed functions, Β55α and B55δ display limited redundancy in this function. During mitosis, chromosomes undergo extensive structural changes, including chromosome condensation, global chromatin compaction, individualization of chromosomes and sister chromatid separation, which allow proper segregation into daughter cells ^39^. Two studies have revealed that Ki-67 is required for the maintenance of spatially separated chromosome arms ^20,40^. The long and highly charged Ki-67 protein promotes the individualization of chromosomes through surfactant-like properties acting as a repulsive layer ^20^. Recent data suggest that during mitotic exit Ki-67 promotes chromosome clustering rather than repulsion, as observed during early mitosis, contributing to the removal of cytoplasm during nuclear assembly ^19^. Our results suggest that both Ki-67 localization and chromosome separation are regulated by PP2A/B55. First, depletion of B55 correlates with increased phosphorylation and accumulation of Ki-67 at mitotic chromosomes in early mitosis. Second, the excessive separation of chromosomes in the scattering phenotype caused by deletion of B55 is suppressed by co-depletion of Ki-67. This is in agreement with a recent report showing that mitotic hyperphophorylation (presumably by CDK1) of the disordered repeat domains of Ki-67 generates alternating charge blocks and promotes their liquid-liquid phase separation (LLPS) to form the periphery of mitotic chromosomes ^41^. Based on all these data together, it is tempting to propose a working model (Figure S5) in which high CDK1 and low PP2A-B55 activities both contribute to promote Ki-67’s function as a surfactant-like protein on the chromosome surface in early phases of mitosis. During mitotic exit, the inactivation of CDK1 and increased activity of PP2A-B55 would lead to the dephosphorylation of Ki-67, reducing its repulsive activity to promote clustering, thereby favouring coordinated chromosome segregation and nuclear assembly during mitotic exit.

## MATERIAL AND METHODS

### Genetically-modified mouse models

The *Ppp2r2a* knockout mouse model was generated by injecting the ES cell clone EPD0328_1_G08 (KOMP Repository) into C57BL/6J blastocysts. Heterozygous *Ppp2r2a* (+/loxfrt) were crossed with EIIa-Cre and pCAG-Flpe transgenic mice to generate the null *Ppp2r2a*(−) and the conditional *Ppp2r2a*(lox) alleles, respectively. The following primers were used for genotyping: *Ppp2r2a_*1F*: 5’-*AAGAATCATGCTGTGCTGCCAAGG-3’; *Ppp2r2a_*1R*: 5’-* CATGCTCTTTATACCTGCCTTATGGACC −3’; *Ppp2r2a_*2R*: 5’-*GGTGCTAGAATTAAGAGTGAGCC-3’. To generate a *Ppp2r2d* knockout model a targeting vector flanking exon 3 of the murine *Ppp2r2d* locus with LoxP sequences was constructed by Gene Bridges, and electroporated in mouse ES cells V6.4 obtained from a hybrid (129 x C57BL/6J) strain. Recombinant ES clones were identified by Southern blot and microinjected into C57BL/6J blastocysts. Heterozygous *Ppp2r2d* (+/loxfrt) were crossed with EIIa-Cre transgenic mice to generate the null *Ppp2r2d* (−) allele. The following primers were used for genotyping: *Ppp2r2d_*1F*: 5’-* GCCACCTGGGGTGTTTTG-3’; *Ppp2r2d_*1R*: 5’-* CATGCTCTTTATACCTTATGGACC-3’.

Mice were maintained on a CD1 x C57BL/6J x 129Sv/J mixed background and were housed in a pathogen-free animal facility at the CNIO following the animal care standards of the institution. The animals were observed on a daily basis, and sick mice were humanely euthanized in accordance with the Guidelines for Humane End Points for Animals used in Biomedical Research (Directive 2010/63/EU of the European Parlament and Council and the Recommendation 2007/526/CE of the European Commission).

### Cell culture, transfection and viral infections

Hela Kyoto cells were kindly provided by Dr. Gerlich ^42^ and mouse embryonic fibroblasts (MEFs) were isolated from E13.5 embryos. Both cell types were maintained in Dulbecco’s modified Eagle’s medium (DMEM) supplemented with 10% foetal bovine serum (FBS) and antibiotics, at 37°C in a humidified 5% CO2 atmosphere. Immortalization of MEFs was performed by transduction with retroviruses expressing the T121 antigen from SV40 virus, selection with hygromycin B (150µg/mL) and passaging for several weeks. Cell synchronization was performed by arresting cells in G0 by serum deprivation (0.1% FBS) during 72 h, followed by addition of complete medium containing 10% FBS serum. When indicated, nocodazole (0.8 µM, Sigma) was added 20h after serum re-addition for the indicated periods of time.

Deletion of *Ppp2r2a* in MEFs was performed by transduction with adenoviruses expressing Cre recombinase (Ad5CMVCre), or adenoviruses expressing Flp recombinase (Ad5CMVFlpo) or empty adenoviruses (Ad5CMVempty) as a control (Viral Vector Core, University of Iowa). Transduction was performed at 250 MOI during 2 days in cells arrested in G0 by serum deprivation. Ki-67 depletion was performed by nucleofection with an siRNA against mouse *Ki-67* (CGUUGAUAUCAGCAACUUU) using Amaxa Nucleofector (Lonza), in accordance with the manufacturer’s instructions.

To knock down *PPP2R2A* and *PPP2R2D* transcripts, specific siRNAs (SI02225825 for *PPP2R2A* and SI02759148 for *PPP2R2D*) were purchased from Qiagen and transfected using Hiperfect (Qiagen), according to manufacturer’s instructions.

pHIV-H2BmRFP was a gift from Bryan Welm & Zena Werb (Addgene plasmid #18982).

### Antibodies and Immunoblotting

The antibody against B55α was generated by immunizing rabbits with a synthetic peptide corresponding to the first 15 amino acids of the N-terminal region of B55α (MAGAGGGNDIQWCFS) conjugated to keyhole limpet hemocyanin (Genscript). The resulting sera were purified with CNBr-activated sepharose 4 Fast Flow (GE Healthcare) conjugated with the same peptide. Purified antibodies were eluted with 0.1M Glycine (pH 2.5), neutralized with 1M Tris pH 8.8 and dialyzed against PBS. PPP2R2D (N2C3) antibody was purchased from Genetex. Ki-67 antibody was purchased from Abcam, and Phospho-Ki-67 antibody was kindly provided by Dr. Imamoto ^24^. β-actin, α-tubulin and vinculin antibodies were all from Sigma.

For immunoblotting, whole-cell extracts were prepared in Laemmli buffer (60 mM Tris pH 6.8; 2% SDS; 10% Glycerol). Proteins were separated on SDS-PAGE, transferred to nitrocellulose membranes and probed with the indicated primary antibodies. HRP-coupled secondary antibodies (DAKO) and ECL system (PerkinElmer) were used for immunodetection.

### Flow Cytometry

Flow cytometry analysis of DNA content was performed by cell fixation with cold 70% Ethanol followed by staining with 10 µg/ml Propidium Iodide (Sigma). Data acquisition was performed with a LSR Fortessa analyzer (BD Biosciences), and FlowJo software was used for data analysis.

### Chromosome spreads

MEFs were exposed to colcemid (0.5µg/mL; KaryoMax) for 6 hours and hypotonically swollen in KCl (75mM) for 30 minutes at 37°C. Cells were then spun down and fixed with Carnoy’s solution (75% pure methanol, 25% glacial acetic acid). After fixation, cells were dropped from a 5-cm height onto glass slides previously treated with 45% of acetic acid. Slides were mounted with Fluoromount-G (SouthernBiotech) and 4,6-diamidino-2-phenylindole (DAPI, Invitrogen) or further processed for immunofluorescence.

### Immunofluorescence

Cells were seeded in coverslips, fixed with 4% formaldehyde for 15 minutes at RT and permeabilized in 0.5% Triton X-100 for 10 minutes at RT. Cells were then blocked with 3% BSA and incubated during 2 hours at RT or ON at 4°C with the indicated primary antibodies, followed by incubation with fluorescent-conjugated secondary antibodies (Molecular Probes, Invitrogen), and staining with DAPI (Invitrogen) to visualize DNA and/or Phalloidin-AF647 (Invitrogen) to visualize actin fibers.

Immunofluorescence over chromosome spreads was performed by rehydrating slides in PBS for 15 minutes, followed by incubation with TEEN buffer (1mM triethanolamine-HCl pH8.5, 0.2 mM NaEDTA, 25mM NaCl) for 7 minutes. Primary and secondary antibodies were diluted in TEEN buffer with 0.5% FBS, and washes were performed in TEEN buffer. Final washing step was done in KB solution (10mM Tris-HCl pH 7.7, 0.15 M NaCl, 0.5% FBS).

Image acquisition was performed using either a Leica D3000 microscope or a TCS-SP5 (AOBS) Laser scanning confocal equipped with an oil immersion objective of 40X (HCX- PLAPO 1.2 N.A.). LASAF v2.6. software was used for image acquisition and analysis was performed with ImageJ or Definiens XD v2.5 softwares, using a customized programmed ruleset.

### Live cell imaging

For videomicroscopy, cells were plated on eight-well glass-bottom dishes (Ibidi) and recorded with a DeltaVision RT imaging system (Olympus IX70/71, Applied Precision) equipped with a Plan Apochromatic 20X/1.42 NA objective lens, and maintained at 37°C in a humidified CO2 chamber. Images were acquired every 7.5 or 10 minutes and analysed using ImageJ software.

### Statistical Analysis

Statistical analysis was carried out using Prism 8 (Graphpad Software Inc.). All statistical tests of comparative data were done using two-sided, unpaired Student’s t-test or ANOVA for differential comparison between two or more groups, respectively. Data with P<0.05 were considered statistically significant (*, P<0,05; **, P<0.01; ***, P<0.001).

## ACKNOWLEDGEMENTS

We thank Dr. Imamoto (RIKEN, Japan) for providing antibodies. M.S.-F. was supported by FPU programme from the Spanish Agencia Estatal de Investigación (AEI) of the Ministry of Science and Innovation (MICIU). M.A.F was supported by the Asociación Española contra el Cáncer (AECC; 2019/INVES19001ALVA). M.R.T was supported by AECC (POSTD211362RUIZ). Research in the laboratory of D.W.G. is supported by the Austrian Academy of Sciences, the Vienna Science and Technology Fund (WWTF; projects LS17-003 and LS19-001), and the European Research Council (ERC) under the European Union’s Horizon 2020 research and innovation programme (grant agreement no. 101019039). Work in the M.M. laboratory was supported by grants from the AEI-MICIIN (RTI2018-095582-B-I00, PID2021-1287260-B-100), the iLUNG2 Programme (S2022/BMD-7437) from the Comunidad de Madrid, and the iDIFFER network of Excellence (RED2022-134792-T). VHIO acknowledges the Cellex Foundation for providing research facilities and equipment and the CERCA Programme from the Generalitat de Catalunya for support. CNIO and VHIO are Severo Ochoa Center<u>s</u> of Excellence (AEI-MICIU CEX2019-000891-S and CEX2020-001024-S/AEI/10.13039/501100011033).

## AUTHOR CONTRIBUTIONS

M.S.-F. performed most experiments, analyzed data and reviewed the manuscript. M.R.-T., C. A.-P., A.E.-B., B.S.-B. and C.V.-B helped with in vitro experiments. S.O. helped in the generation of the mouse models and D.M. contributed to microscope analysis. D.W.G. provided reagents, analyzed data and reviewed the manuscript. M.A.-F. and M.M. analyzed data, supervised the project and wrote the manuscript.

