## Supplemental Information for "PP2A-B55α,δ phosphatase counteracts Ki67-dependent chromosome individualization during mitosis"

Sanz-Flores et al.

### SUPPLEMENTARY MATERIAL

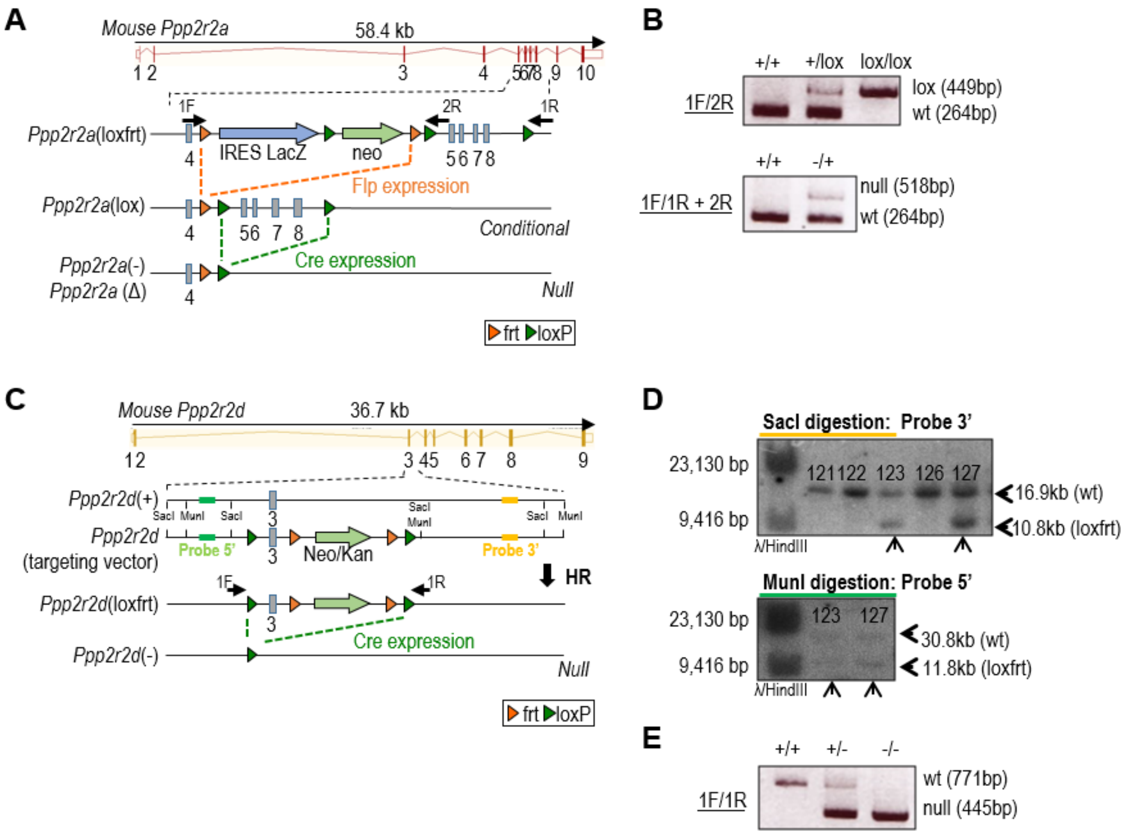


**Figure S1. Generation of *Ppp2r2a* and *Ppp2r2d* knockout mouse models. A,** Schematic representation of the *Ppp2r2a* genomic structure and the conditional knockout allele used in this study. LoxP (green triangles) sites and frt (orange triangles) sites were used to flank *Ppp2r2a* exons 5-8 and the neo-resistance and IRES-LacZ cassettes, respectively. Flp-mediated recombination results in the *Ppp2r2a*(lox) allele. Site-specific Cre recombination results in the germline *Ppp2r2a*(–) or the conditionally-induced *Ppp2r2a*(Δ) null alleles **B,** Representative PCR products showing the indicated *Ppp2r2a* alleles after amplification from tail genomic DNA, using oligonucleotides indicated in A. **C,** Schematic representation of the *Ppp2r2d* genomic structure and the knockout allele used in this work. LoxP (green triangles) sites are used to flank exon 3 of *Ppp2r2d,* and the neo-resistance cassette, which has been used to select positive clones for homologous recombination (HR) in ES cells. Site-specific Cre recombination results in the germline *Ppp2r2d*(–) null allele used in this study. **D,** Southern blot analysis of recombinant ES cells showing two *Ppp2r2d*(+/loxfrt) positive clones (marked with arrows) that underwent HR. Position of 5’ and 3’ probes and restriction enzymes used for Southern blot analysis are indicated in C. Positive clones, detected upon hybridization with the 3’ probe, were then hybridized with the 5’ probe for confirmation. **E,** Representative PCR products showing the indicated *Ppp2r2d* alleles after amplification from tail genomic DNA, using oligonucleotides indicated in C.

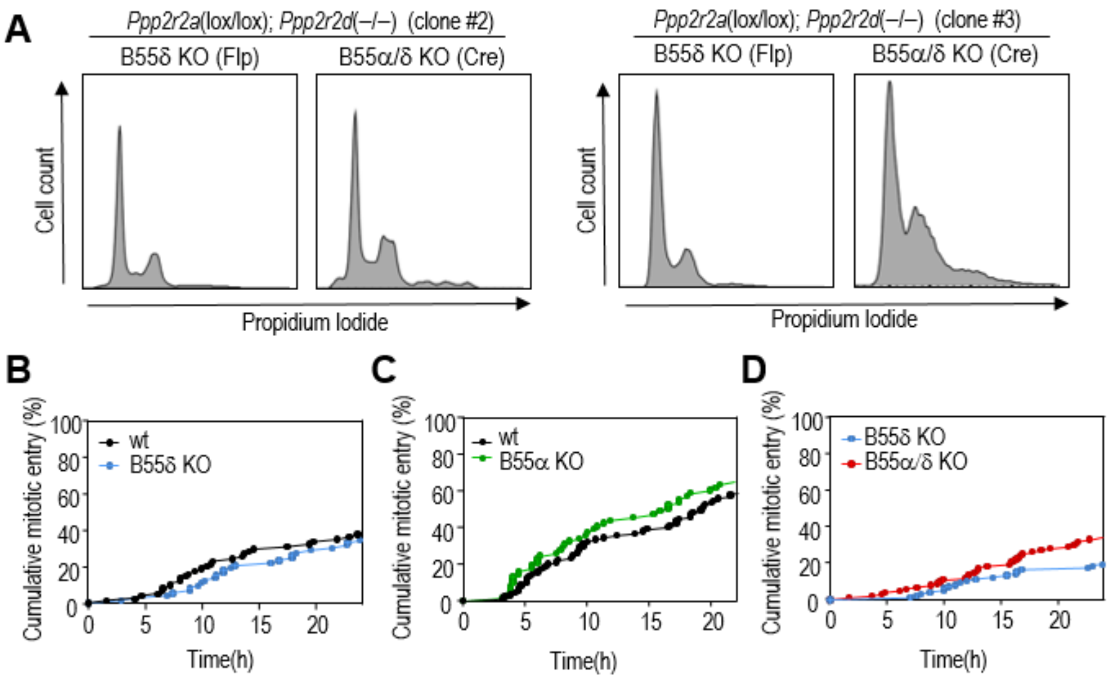


**Figure S2. Cell cycle analysis and mitotic entry in B55α/δ KO cells. A,** Cell cycle profile (propidium iodide staining) of two *Ppp2r2a*(lox/lox); *Ppp2r2d*(–/–) clones 6 days after infection with AdenoFlp (B55δ KO) or AdenoCre (B55α/δ KO) viruses. **B-D,** Videomicroscopy analysis of mitotic entry (scored by cell rounding and chromosome condensation) in B55δ KO **(B)**, B55α KO **(C)** and double B55α/δ KO **(D)** cells in comparison with their respective controls. One representative clone per genotype is shown.

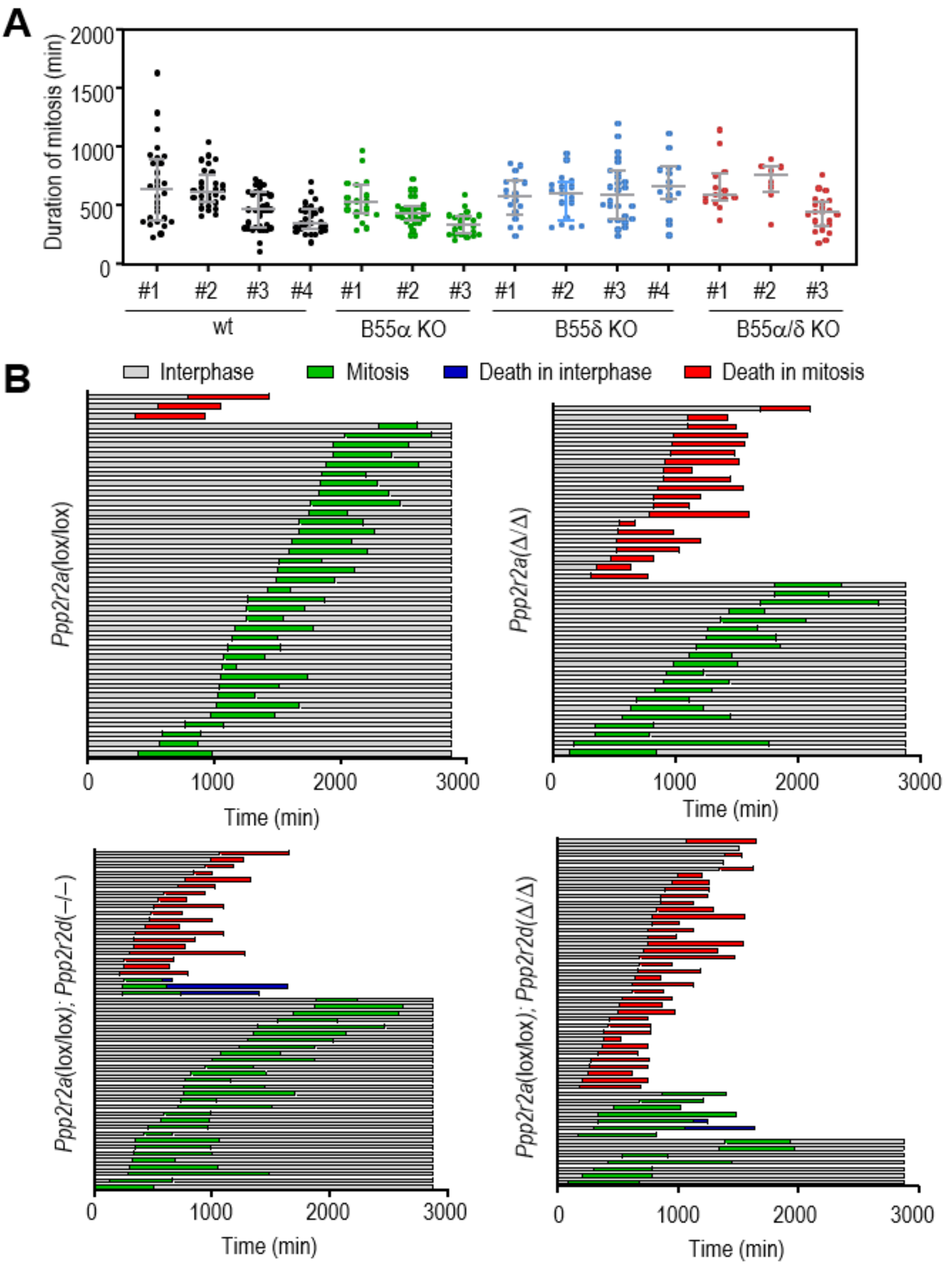


**Figure S3. Analysis of mitosis in B55 deficient cells in the presence of nocodazole. A,** Duration of mitosis (DOM) from NEB until chromosome decondensation in wt, B55α KO, B55δ KO and B55α/δ KO iMEFs in the presence of nocodazoles (0.8 µM). Data are means ± SD of each clone (n=3-4 clones per genotype). **B,** Cell fate analysis by time-lapse microscopy of cells with the indicated genotypes upon nocodazole treatment. Each row represents an individual cell.

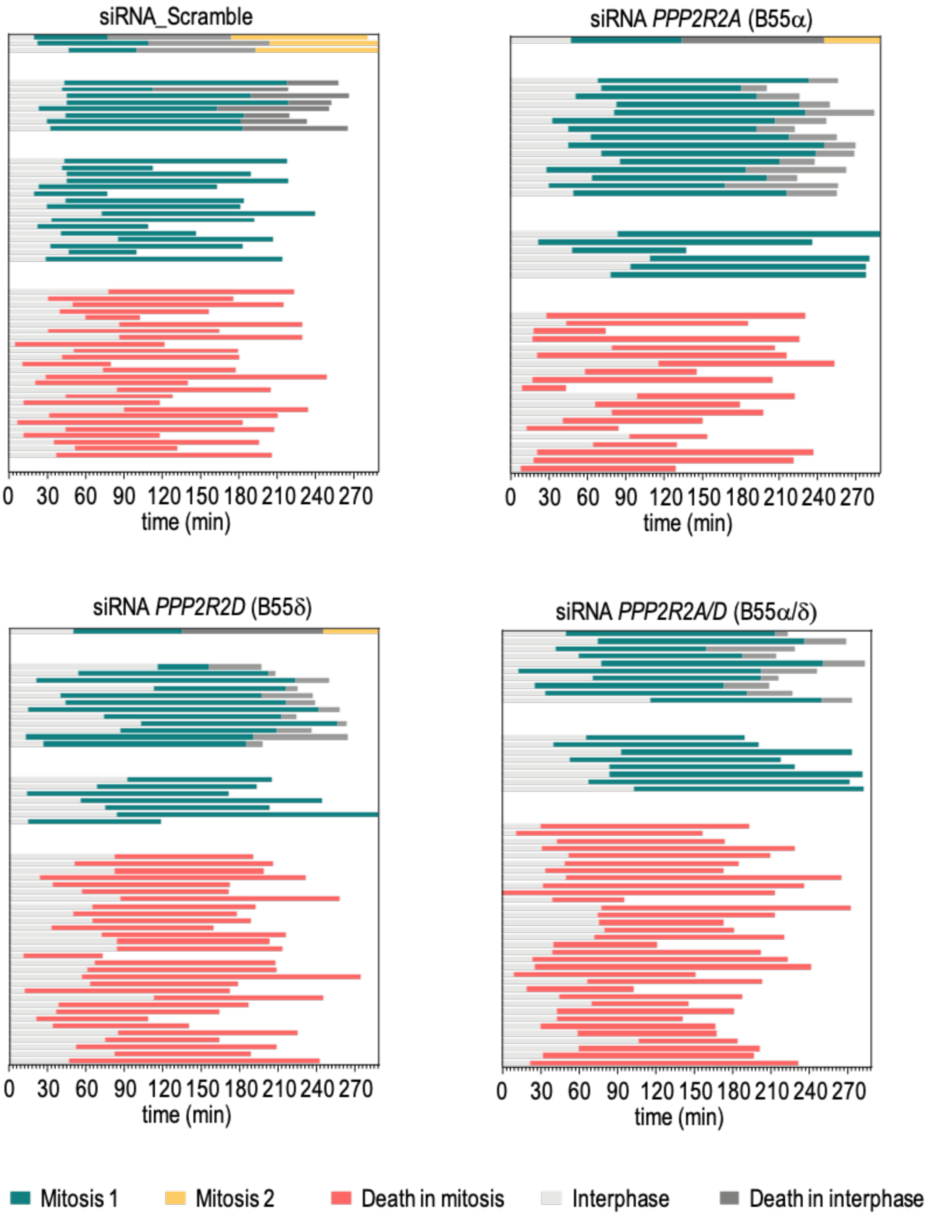


**Figure S4. Analysis of mitosis in B55 depleted Hela cells in the presence of nocodazole.** Cell fate analysis by time-lapse microscopy of Hela cells transfected with the indicated siRNAs upon nocodazole treatment. Each row represents an individual cell.

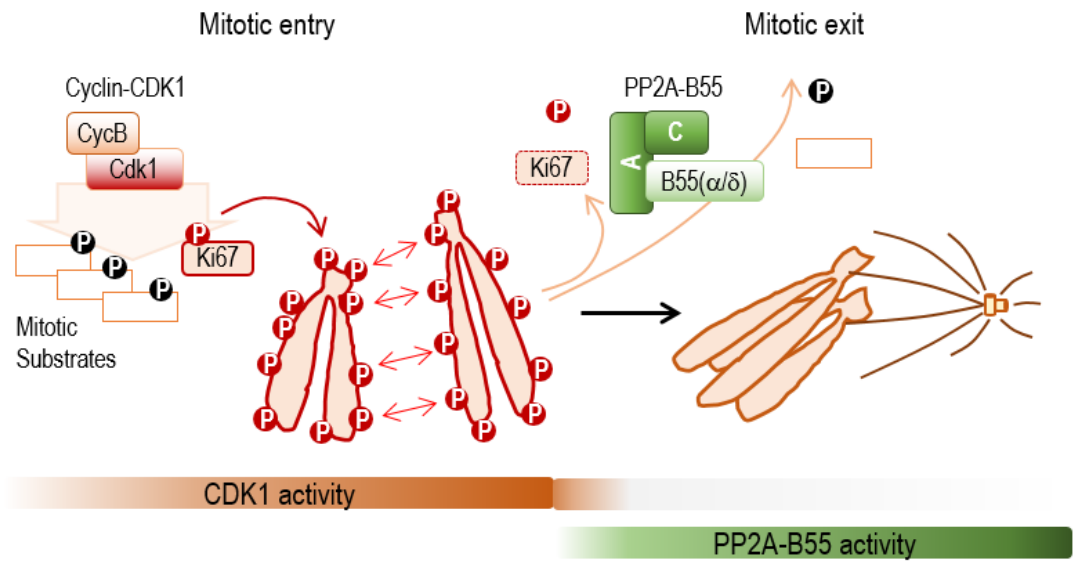


**Figure S5. A model for the role of PP2A-B55 complexes in chromosome clustering.** Ki-67 is phosphorylated at least partially by CDK1 during mitotic entry promoting its accumulation at the chromosomal periphery and thereby resulting in chromosomal individualization, a process required for proper movements and bipolar attachment of chromosome during the early phases of mitosis. Dephosphorylation of Ki-67 by PP2A-B55$\alpha$,$\delta$ complexes allow clustering of chromosomes and likely favours, together with microtubule attachment, the coordinated movement of the chromosomal mass towards the division poles.
